# Computational joint action: dynamical models to understand the development of joint coordination

**DOI:** 10.1101/2024.02.25.582011

**Authors:** Cecilia De Vicariis, Vinil T. Chackochan, Laura Bandini, Eleonora Ravaschio, Vittorio Sanguineti

## Abstract

Coordinating with others is part of our everyday experience. Previous studies using sensorimotor coordination games suggest that human dyads develop coordination strategies that can be interpreted as Nash equilibria. However, if the players are uncertain about what their partner is doing, they develop coordination strategies which are robust to the actual partner’s actions. This has suggested that humans select their actions based on an explicit prediction of what the partner will be doing – a partner model – which is probabilistic by nature. However, the mechanisms underlying the development of a joint coordination over repeated trials remain unknown. Very much like sensorimotor adaptation of individuals to external perturbations (eg force fields or visual rotations), dynamical models may help to understand how joint coordination develops over repeated trials.

Here we present a general computational model – based on game theory and Bayesian estimation – designed to understand the mechanisms underlying the development of a joint coordination over repeated trials. Joint tasks are modeled as quadratic games, where each participant’s task is expressed as a quadratic cost function. Each participant predicts their partner’s next move (partner model) by optimally combining predictions and sensory observations, and selects their actions through a stochastic optimization of its expected cost, given the partner model. The model parameters include perceptual uncertainty (sensory noise), partner representation (retention rate and process noise), uncertainty in action selection and its rate of decay (which can be interpreted as the action’s learning rate). The model can be used in two ways: (i) to simulate interactive behaviors, thus helping to make specific predictions in the context of a given joint action scenario; and (ii) to analyze the action time series in actual experiments, thus providing quantitative metrics that describe individual behaviors during an actual joint action.

We demonstrate the model in a variety of joint action scenarios. In a sensorimotor version of the Stag Hunt game, the model predicts that different representations of the partner lead to different Nash equilibria. In a joint two via-point (2-VP) reaching task, in which the actions consist of complex trajectories, the model captures well the observed temporal evolution of performance. For this task we also estimated the model parameters from experimental observations, which provided a comprehensive characterization of individual dyad participants.

Computational models of joint action may help identifying the factors preventing or facilitating the development of coordination. They can be used in clinical settings, to interpret the observed behaviors in individuals with impaired interaction capabilities. They may also provide a theoretical basis to devise artificial agents that establish forms of coordination that facilitate neuromotor recovery.

**Author summary:** Acting together (joint action) is part of everyday experience. But, how do we learn to coordinate with others and collaborate? Using a combination of experiments and computational models we show that through multiple repetitions of the same joint task we select the action which represents the ‘best response’ to what we believe our opponent will do. Such a belief about our partner (partner model) is developed gradually, by optimally combining prior assumptions (how repeatable or how erratic our opponent behaves) with sensory information about our opponent’s past actions. Rooted in game theory and Bayesian estimation, the model accounts for the development of the mutual ‘trust’ among partners which is essential for establishing a mutually advantageous collaboration, and explains how we combine decisions and movements in complex coordination scenarios. The model can be used as a generative tool, to simulate the development of coordination in a specific joint action scenario, and as an analytic tool to characterize the individual traits or defects in the ability to establish collaborations.

## Introduction

Many of our everyday activities take place in social settings and are coordinated with other people. Even seemingly simple interactions, like a pair of workers sawing timber with back and forth movements, a couple executing complex steps on a dance floor, two children playing badminton, or providing physical therapy to a patient, require that two individual minds are somehow connected and their bodies coordinated [1]. The characteristic feature of these interactions is that participants influence each others’ behavior through coupled sensorimotor exchanges within continuous action spaces, continuously in time, and possibly over repeated trials. Many of these interactions require active coordination, which is manifested by physical and cognitive responses related to explicit knowledge of the interacting partner.

When the two humans are physically coupled through a tool [2] or a pair of robot devices [3], haptics is a major source of information about their partner. These scenarios have been particularly useful for quantifying various aspects of joint action performance, because dual haptic or ‘dyadic’ interfaces can be programmed to implement a variety of interaction modalities and allow to experimentally manipulate all aspects of the interaction [3–6].

During the last two decades, discrete-time dynamical models have been used to investigate sensorimotor learning and adaptation. These models have helped to clarify the underlying mechanisms: the role of the sensory prediction error [7, 8]; the interplay of implicit and explicit adaptation mechanisms [9]; and the evidence of multiple concurrent adaptation processes across different time scales [10]. The same modeling framework has been used to assess how adaptation is altered in a number of neurological conditions, such as cerebellar degeneration [11] and multiple sclerosis [12], and to model neuromotor recovery through robot-assisted exercise [13].

A similar approach can be used to gain more insight on how we develop coordination strategies while we interact with our peers. Few studies have addressed the perceptual and control mechanisms of joint action [4–6, 14]. Dynamical models rely on optimality principles to account for perception and action selection within a single trial. Perception is believed to optimally combine prior beliefs and the available observations to minimize the prediction uncertainty [15, 16]. These same Bayesian mechanisms likely mediate the prediction of the actions of our partners [6, 14]. Partial information about the partner likely affects the development of a joint coordination. Action selection in joint coordination can be interpreted in terms of game theory. Several studies [4, 6, 17] have reported that the sensorimotor behavior of physically coupled subjects converges toward a Nash equilibrium – a situation in which none of the players can unilaterally improve their payoff [18]. However, the vast majority of previous studies addressing joint action within a game theoretic framework only focus on equilibrium situations [4, 17]. Very few studies [6, 19, 20] have addressed the way joint coordination is negotiated and learned in scenarios that involve movements. One simple learning strategy when the players play repeatedly the game is that at every round each player determines their best response based on their beliefs about how their opponents will play (fictitious play, FP) – see [21, 22]. FP has been implied as a candidate mechanism for the development of joint decision-making [19] or joint action in humans [6], at least in situations where the players use stationary strategies. Reinforcement learning mechanisms have been also implicated in situations where the players have knowledge about their task [20].

Here we present a general modeling framework to describe the development of coordination strategies in joint action. The model addresses a variety of interactive scenarios and can be used both as a simulation and an analytical tool. We first demonstrate the model with a toy problem – a sensorimotor version of a classical game with multiple Nash equilibria (Stag Hunt). We then focus on joint sensorimotor coordination games [6], in which participants are mechanically coupled, have a shared final target, and are instructed to cross a via-point (VP, different for each player) while minimizing their interaction force. We estimate the model parameters from the experimental data (action time series over trials). We show that the model correctly captures the different experimental conditions and correctly reproduces, for each condition, the temporal evolution of the coordination strategies.

## Model

We specifically focus on situations in which the players know their own goal, but not their partner’s, and they have imperfect information about their partner’s action. We also assume that the agents develop a coordination over multiple rounds of the same game. Many joint action studies fall within this scenario. We will later discuss how these assumptions can be relaxed in order to deal with more general situations.

The overall model is depicted in Figure 1. At each trial *t*, the *i*-th player selects their own action, *u*_*i*_(*t*), on the basis of the partner’s expected move (partner model), *x*_*i*_(*t*) = û _−*i*_(*t*) – we use −*i* to indicate the player other than *i*. Once the actions have been carried out, through their sensory systems both players collect information *y*_*i*_(*t*) about their own and their partner’s actions. This information is then used to update their partner model, which will be accounted for when selecting the action on the next trial.

**Fig 1.**
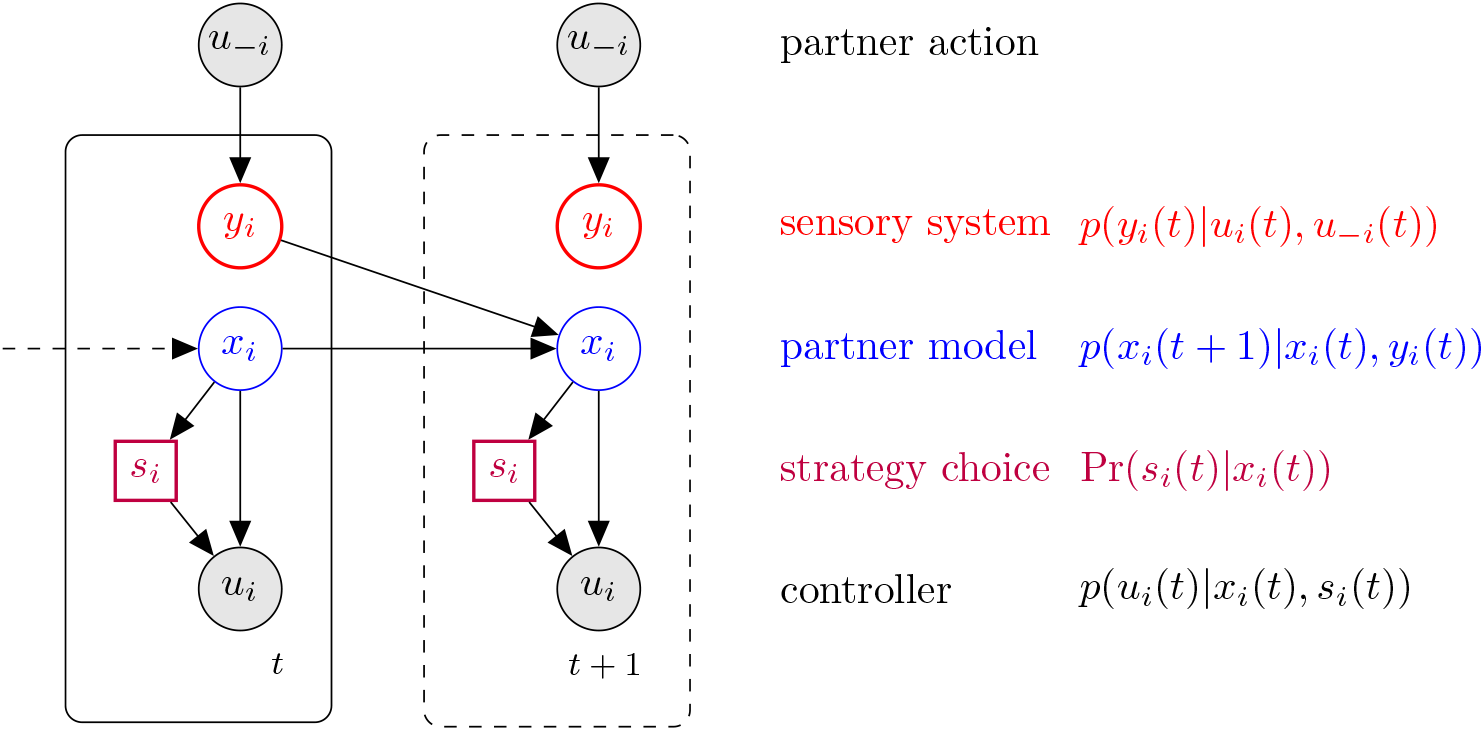
Probabilistic model of a single joint action player, that to account for multiple strategy choices. At each time step, Player *i* receives information about partner action (*u*_−*i*_) through their sensory system (*y*_*i*_). This information is used to predict the next partner action (partner model, *x*_*i*_). The partner model is used to select the actual action (*u*_*i*_) which also depends on the strategy choice (*s*_*i*_). The latter decision is also affected by the player’s knowledge about the partner. For each player, sensory input (*y*_*i*_), partner model (*x*_*i*_) and strategy choice (*s*_*i*_) are latent variables; actions (*u*_*i*_ and *u*_−*i*_) are observable variables (denoted by shaded nodes)

### Joint tasks as mixtures of quadratic games

Individual motor tasks are often described in terms of a quadratic cost function which accounts for the trade-off between error and effort [24]. For the *i*-th player, the model specifies the task as a quadratic cost function, which depends on both players’ actions *u*_1_ and *u*_2_:

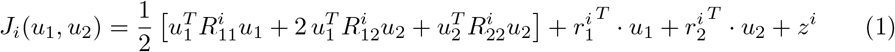

Static games with infinite action space and quadratic costs are known as quadratic games [23]. The quadratic game formalism naturally extends individual motor control to joint action scenarios.

If all 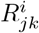 are symmetric and positive definite, a quadratic game has a single Nash equilibrium [23] which is calculated as:

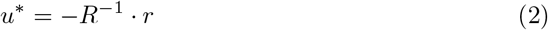

where 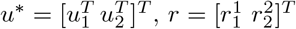 and:

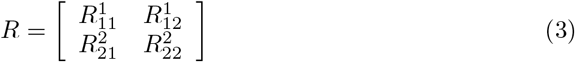

Many joint action scenarios involve selecting an action among a set of alternatives – for instance, reaching one among several possible targets. To extend the quadratic game scenario to these situations, we assume that the cost incurred by player *i* depends on a decision variable 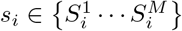, which specifies the decision between one of the *M* options that the *i*-th player has available. Therefore, Eq. 1 becomes:

### Sensory system

At every trial *t*, the sensory system of each player provides an estimate of their partner’s action:

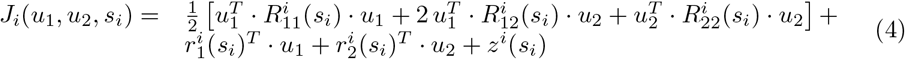

where 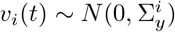 is the sensory (measurement) noise of the *i*-th player, with covariance 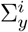. Parameter *H*^*i*^ reflects the structure of the sensory system and relates partner action to its sensory consequences.

### Partner model

We assume that at every trial, *t*, each player predicts their partner’s next action by optimally combining predictions and actual observations – provided by the sensory system above – into a ‘partner model’. The *i*-th player’s prior belief about partner action *u*_−*i*_ can be summarized as:

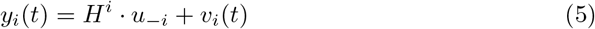

where 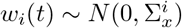 is a process noise and *A*^*i*^ is a retention (memory) parameter. Parameters *A*^*i*^ and 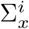 reflect the player’s prior belief on partner behavior – respectively, the extent to which partner behavior is stationary, or is erratic across trials. In particular, *A*^*i*^ = 0 denotes a memoryless partner behavior and 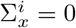 denotes a purely deterministic partner behavior.

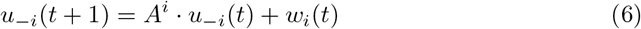

The Bayes-optimal solution to this integration problem is provided by the Kalman filter algorithm. Here we only provide the final formulation, complete derivation is provided in the Supplementary Note. At each time step *t* the partner model of the *i*-th player provides a prior estimate (prediction) of partner action, defined by its mean *x*_*i*_(*t*) and covariance *P*_*i*_(*t*). These quantities are calculated by incorporating the sensory outcome of the actions performed at the previous iteration:

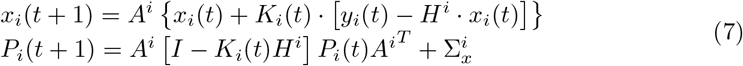

where the Kalman gain 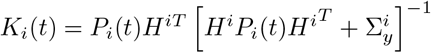 is calculated iteratively. The portion in {… } in Equation 7 optimally combines the action prediction at time *t, x*_*i*_(*t*) with the sensory information obtained after both players have performed their actions. Parameter *A*^*i*^ determines the propagation of this estimate to the predicted action at the next step.

### Action selection and strategy choice

The optimal action of the *i*-th player at time *t, u*_*i*_(*t*) – and the optimal strategy *s*_*i*_(*t*) if multiple options exist – minimizes the expected cost 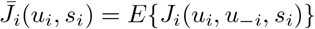, where the expectation is calculated over the partner’s action, *u*_−*i*_ (actually, player’s belief about partner actions). At each trial the partner model provides the expected value *x*_*i*_(*t*) of the partner’s action and its covariance matrix *P*_*i*_(*t*), so that the expected cost can be expressed in terms of these quantities. The expected cost turns out to be a quadratic form in *u*_*i*_:

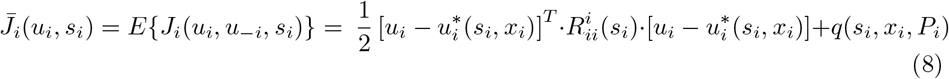

whose parameters depend on *x*_*i*_ and *s*_*i*_ – see the Supplementary Note for complete derivation.

To account for variability in action selection – i.e. players not behaving in a deterministic way, or having an imperfect knowledge of their cost function – we assume that the selection of both strategy and action is in fact a stochastic process. The expected cost can be interpreted as an energy term so that the joint probability of *u*_*i*_ and *s*_*i*_ given the partner model can be expressed as a Boltzmann probability density function:

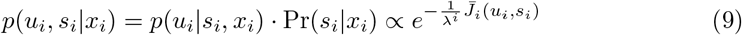

where *λ*^*i*^ reflects action selection ‘temperature’, with greater *λ*^*i*^ leading to greater randomness.

In the special case of 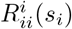 being all positive definite, the action probability given the partner model *x*_*i*_ and the strategy 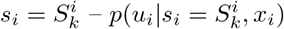 has a multivariate Gaussian distribution, with mean 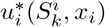 and covariance 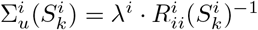. The probability of selecting strategy 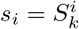 given the partner model *x*_*i*_ also depends on partner model and game structure:

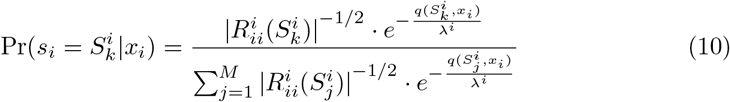

See the Supplementary Note for complete derivation. As a consequence, when the 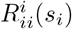 are all positive definite, the action *u*_*i*_ given the partner model *x*_*i*_ has a Gaussian Mixture distribution.

The proposed action generation mechanism naturally satisfies the minimum intervention principle [25]: in a redundant task, the action cost does not depend on the task-irrelevant movement features and, conversely, is very sensitive to task-relevant features, so that action selection variability is modulated by task relevance. Further, the action generation model can be seen as an extension of the quantal response equilibria (QRE) model [26] to continuous games. As in the QRE, strategy choice depends on the task and is modulated by partner model uncertainty – the dependence of *q*(*s*_*i*_, *x*_*i*_) on *P*_*i*_. Likewise, probabilistic action selection reflects the task requirements and is modulated by the temperature parameter. In many learning scenarios, the selected actions are expected to gradually become more repeatable from trial to trial. To allow the ‘temperature’ parameter *λ*^*i*^(*t*) to decrease over trials, in the model we specify the initial temperature, 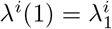, and a decay rate *a*^*i*^ < 1, so that

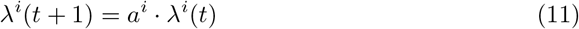

In the above derivation, we made the critical assumption that the 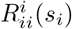 are positive definite. In this case, the probability distribution of actions – is a Gaussian or a Gaussian Mixture. However, this condition may not be met in many types of scenarios, e.g. competitive games where the gain of one player corresponds to the loss of the other; in this situation the matrix 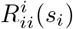 may no longer be positive definite for some of the players. Nevertheless, if the action space is limited, Eq. 9 still defines a valid action probability density function, though no longer a mixture of Gaussians.

At every round, each agent determines their best response to the empirical distribution of their partner’s action. This process, known as fictitious play (FP) [22], is in many cases sufficient to learn a coordination strategy.

One player’s behavior is specified in terms of seven parameters. The sensory noise covariance 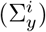 specifies the reliability of the player’s sensory system. The behavior of the partner model is specified by the initial partner model mean (*μ*^*i*^) and covariance 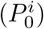, whereas retention rate (*A*^*i*^) and process noise 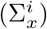 specify the way it evolves with time. Taken together, these parameters specify the player’s prior information about the partner. Over trials, this information is optimally combined with the sensory information. Finally, initial temperature 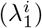 and temperature decay rate (*a*^*i*^) specify the degree of randomness in action selection and how it evolves with time.

The model is general enough to simulate the trial-by-trial evolution of a large variety of joint action scenarios which involve perception and action. Further, the model is simple enough to be used to analyze joint action experiments. Given a time series of measured players’ joint actions, it is possible to estimate the model parameters that best describe each player in the dyad. The Supplementary Note describes a maximum likelihood procedure to estimate the model parameters.

## Results

We demonstrate the model in a variety of interaction scenarios, in which two players engage in a sequence of interactions. We specifically focus on how convergence to a joint strategy is determined by the information available about their partner. In all scenarios, we show that the model generates plausible action sequences. When experimental results are available, they are compared with the model predictions.

### Spatial Stag Hunt

We first demonstrate the model behavior in a toy (simulated) experiment, inspired by the classical Stag Hunt game. Two hunters have the option of jointly hunting a stag (S), or independently pursuing a rabbit (R). Hunting a stag results in a greater payoff, but can only be done if both partners cooperate. Catching a rabbit does not require partner cooperation, but results in a smaller payoff. This game has two Nash equilibria: both hunters catching a rabbit (R-R), and both hunters catching the stag (S-S). The R-R strategy has a lower payoff than the S-S strategy. The original version of the game assumes a discrete action space (R,S). Yoshida et al. (2008) [27] formulated a spatial version of the game, in which two players are required to move their hand within a one-dimensional workspace toward either a rabbit of a stag, located at fixed positions *u*_*R*_ and *u*_*S*_. The participants are supposed to act simultaneously, without knowing the decision of the partner. The payoff for catching the stag can only be achieved when both players are in *u*_*S*_, whereas the rabbit payoff does not depend on the other agent’s action.

Our modeling framework explicitly focuses on joint actions that involve movements and perception. This requires some extra assumptions about the physical arrangement of the simulated experiment. Specifically, we assumed that at each trial each player controls a cursor along a line representing the action space. The locations of both the rabbit and stag are fixed and known to both players. We additionally assumed that at the end of each trial, each player is provided with imperfect information on their partner’s location. In a physical experiment, this could be done by a blurred display (random dots). The above assumptions are summarized in Figure 2 (left).

**Fig 2.**
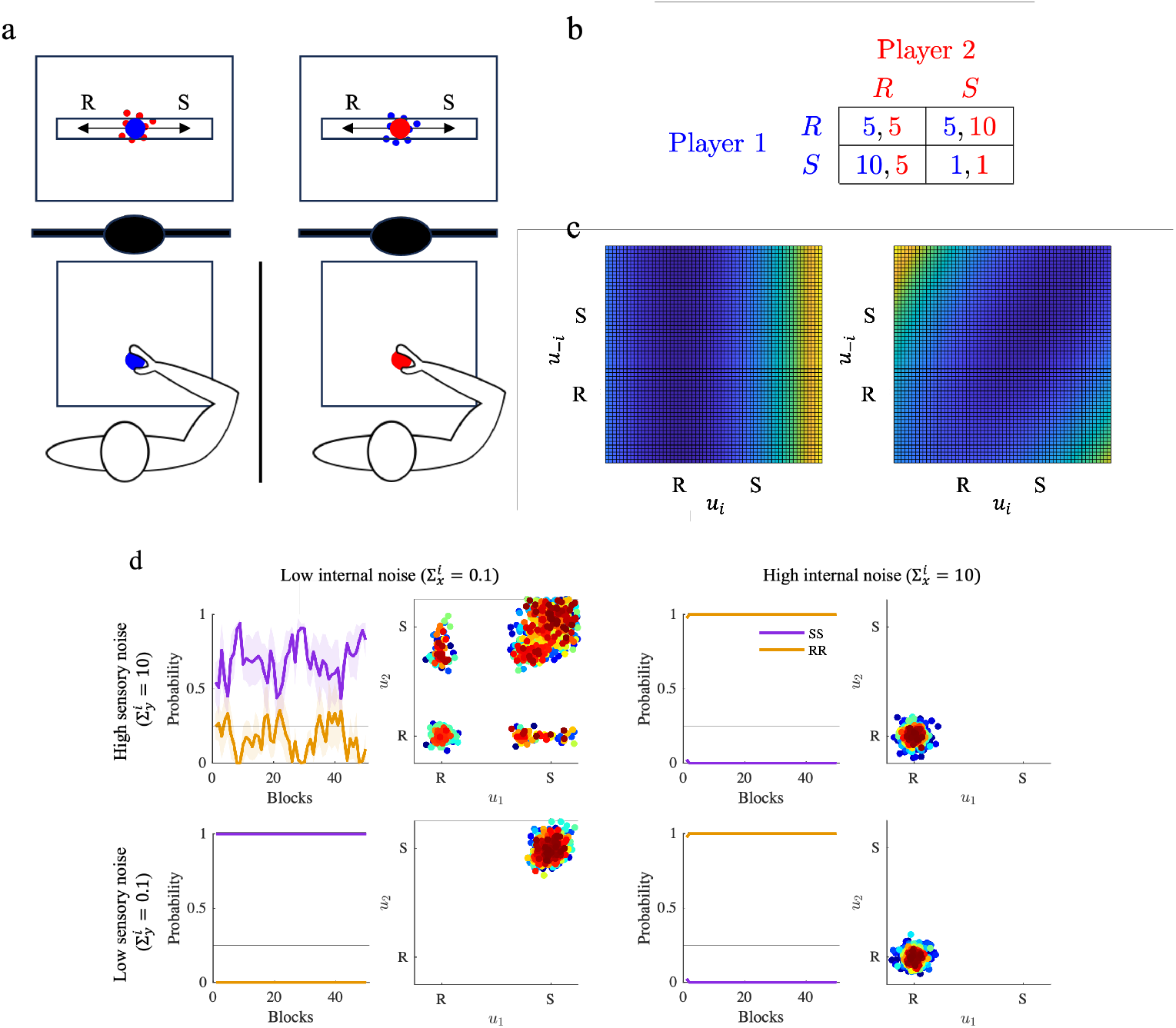
Stag Hunt game. a. simulation concept. Each player knows the locations of both rabbit (R) and stag (S). They can see their actions and have imperfect information about their partner’s actions (blurred circles). b. cost matrix used in the simulations, in which the cost can be interpreted as a reduced payoff. c. Costs related to the two equilibria are represented in the action space. Cold colors indicate lower costs, and hot colors indicate higher costs. On the left, whichever action the partner selects it is safe for player *i* to go for the Rabbit solution. On the right, the two players minimize their costs by selecting both the Stag. d. Simulated Stag Hunt game for different combinations of sensory noise 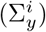 and partner predictability (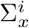 . Temporal evolution of the probabilities of selecting the SS (blue), the RR (orange)nThe curves represent game performance at population level, where probabilities (mean *±* SE) have been computed over epochs (1 epoch = 40 trials) and averaged over multiple simulated dyads. The horizontal line indicates the chance level (p=0.25). Bottom: scatter plots of the joint actions (*u*_1_, *u*_2_) for one representative dyad for each condition

To account for the two Nash equilibria of this game, we introduce a discrete variable *s*_*i*_ which denotes whether player *i* is pursuing the ‘rabbit’ (*s*_*i*_ = 1) or the ‘stag’ (*s*_*i*_ = 0). Therefore, the cost function for player *i* is:

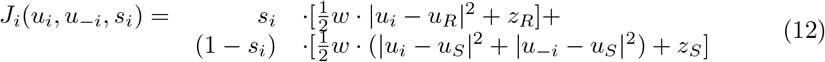

Details on the calculation of the model parameters on the basis of the cost matrix of Figure 2 (right) are provided in the Methods section. We additionally assume that at the end of each trial, both participants are displayed the cost incurred according to Eq. 12. Both are instructed to keep their cost as low as possible.

We simulated the model by varying the reliability of sensory information (‘low’ or ‘high’ 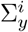) and the belief of partner predictability (‘low or ‘high’ 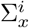). The simulation results are summarized in Figure 2. Convergence to the S-S strategy critically requires that the partner is believed to be predictable (low 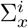), and reliable sensory information (low 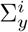) about the partner. Conversely, when partner predictability is low, irrespective of 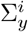 the R-R strategy is preferred. When sensory information is less reliable, the players’ behaviors become more erratic and tend to fluctuate between SS and RR. These observations are confirmed by statistical analysis. We used a two-way ANOVA to compare the four groups of simulations. We used sensory and internal noise as factors. We found a significant effect of sensory noise – *F* (28, 1) = 98.88, *p* <0.001 – and an even stronger effect of internal noise – *F* (28, 1) = 2428.63 *p* <0.001. The sensory and internal noise interaction is also significant – *F* (28, 1) = 98.88, *p* <0.001.

The model suggests that convergence to the S-S Nash equilibrium requires two conditions: (i) reliable sensory information (low 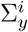) and (ii) a belief in partner predictability 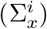. If the second condition is not satisfied, the partner model would only rely on sensory information, the actions would be purely reactive. This would prevent the build-up of trust which is necessary to establish a robust collaboration.

As a cross-check, we identified the model parameters from the simulated actions time series. To robustly recover model parameters, the action time series must exhibit some temporal variability, a situation which only occurred in the low-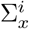 and high-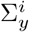 condition; see Figure 2d. For this condition we indeed obtained a high correlation coefficient (*R*^2^ *>* 0.95) for the estimated actions time series and small absolute errors between the simulated and estimated model parameters - details are reported in the Supplementary File.

### Two Via-Point game

In the two via-point (2-VP) task [6], the players are mechanically connected through a virtual spring. They perform reaching movements from the same start and end position, by crossing a via-point (VP) which is different for the two players. Each player can only see their own VP, but not their partner’s. Both are instructed to keep the spring force – and therefore the inter-player distance – as low as possible during movement; see Figure 3a. The joint task has two Nash equilibria, in which both players follow the same path and cross both via-points in the same order, either VP_1_ → VP_2_ or VP_2_ → VP_1_. Depending on the VP locations, the two Nash equilibria may substantially differ in terms of path length and effort required. When the VPs are arranged along the midline of the start-end segment, the two Nash equilibria have exactly the same cost – see Figure 3b.

**Fig 3.**
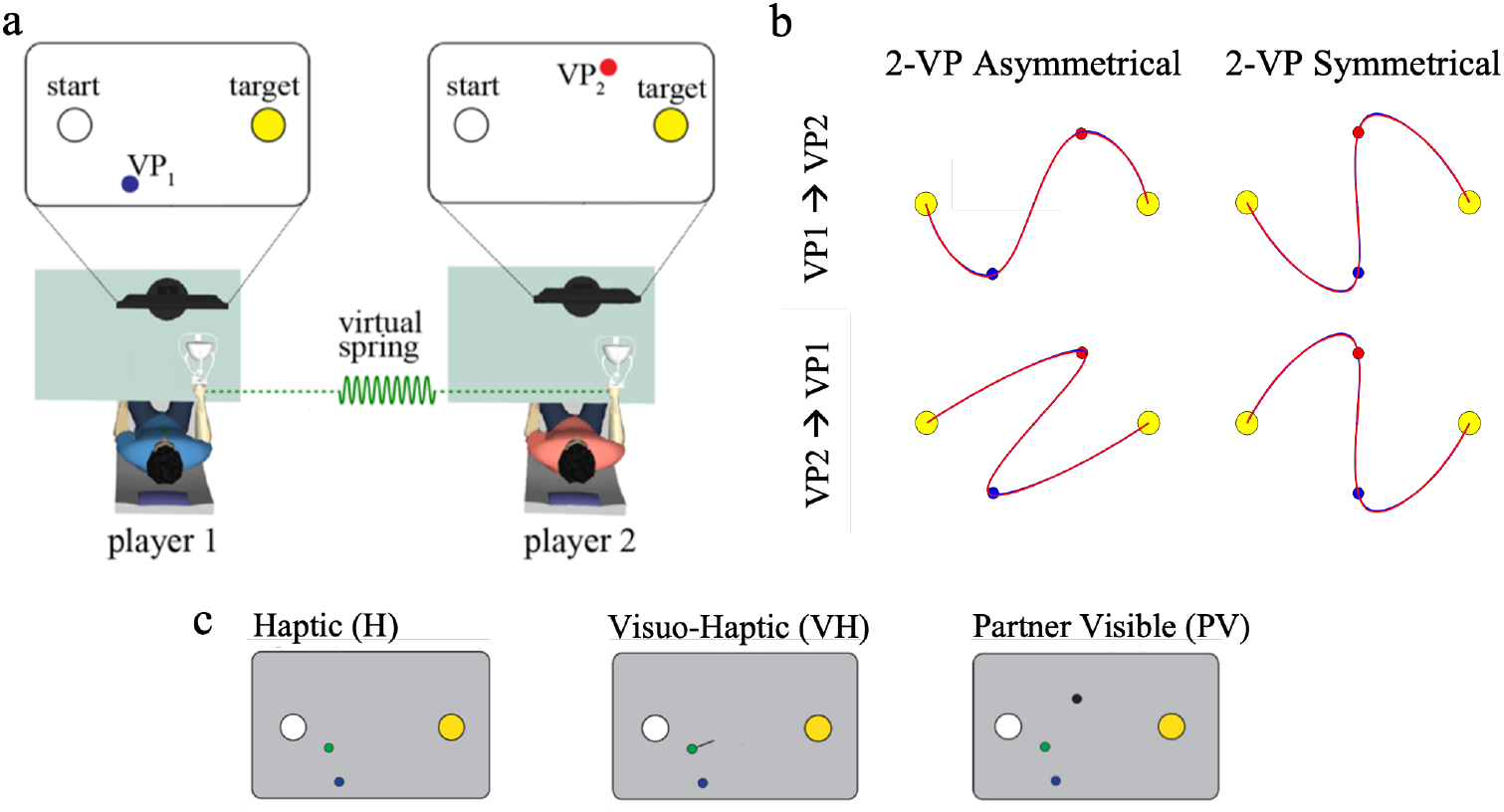
The 2-VP task. a. Experimental apparatus and task – reprinted from [6]. The partners were connected through a virtual spring. b. Nash equilibria with asymmetric and symmetric via-points locations. c. In different experimental groups, information about partner actions was provided haptically, through the interaction force alone (Haptic group, H); by additionally displaying the interaction force vector on the screen (Visuo-Haptic group, VH), or by also displaying the partner’s cursor (Partner Visible, PV).

In terms of the model, the 2-VP task can be described by the following cost function (one per player):

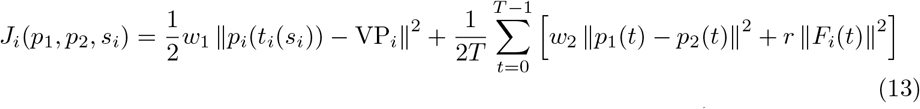

where *t* = 0 is taken as the start time and *t* = *T* as the final time (common to both players) and we assume that *p*_*i*_(0) = *p*_*start*_ and *p*_*i*_(*T*) = *p*_*end*_. Time *t*_*i*_ is the time at which the *i*-th player crosses their own via-point VP_*i*_. The crossing time is determined by the decision variable *s*_*i*_, which specifies whether the VP is crossed early (*s*_*i*_ = *E*), in the middle (*s*_*i*_ = *M*), or late in the movement (*s*_*i*_ = *L*). We make the simplifying assumption that, across trials, the crossing time *t*_*i*_(*s*_*i*_) only takes three fixed values. Body dynamics of the *i*-th player can be described as

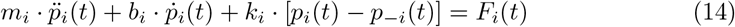

where *F*_*i*_(*t*) is the motor command.

In [6] the task was modeled using differential game theory, and the resulting strategies were specified by a pair of feedback controllers. Given the short duration of one trial, here we make the simplifying assumption that the action of one player only affects their partner’s in future trials, whereas within-trial corrections are neglectable. Under this assumption, the proposed modeling framework is appropriate to describe this task. We just need a parametric representation of the player’s action within a single trial. We approximate the whole trajectory *p*_*i*_(·) for each player with a polyharmonic spline, specified by a *M* nodes *u*_*i*_ = [*p*_*i*_(*t*_1_) … *p*_*i*_(*t*_*M*_)]^*T*^ . We define the ‘action’ of the *i*-th player at a given trial as the vector of nodes’ coordinates, *u*_*i*_; see the Methods section for details. This approximation captures the overall shape of a generic trajectory in terms of the coordinates of a small number of nodes. Subsequent derivatives of *p*_*i*_(*t*) (velocity, acceleration) can also be expressed as linear combinations of the *u*_*i*_’s.

In different experimental groups, in a previous study [6] we varied the information provided about the ongoing partner actions, namely haptically, through the interaction force alone (Haptic group, H); by additionally displaying the interaction force vector on the screen (Visuo-Haptic group, VH) or by also displaying the partner’s cursor (Partner Visible, PV); see Figure 3c. For the first two experimental conditions (H and VH), the players see their own hand and feel the interaction force, which depends on the locations of both hands. In the PV condition, the position of their partner’s hand is also displayed on the screen. Therefore, all groups have equal information on their own hand (vision and haptics), whereas the uncertainty in perceiving their partner is minimal in group PV and maximum in group H.

#### Experiment 1: asymmetric via-points

Looking at the experimental findings, all participants developed strategies that are consistent with the low-effort Nash equilibrium (crossing VP_1_ first, then VP_2_), corresponding to *s*_1_ = *E* and *s*_2_ = *L*. The hand trajectories turned out to be increasingly close to the Nash equilibrium when information about the partner was more reliable (PV group). In the model, we set *t*_*i*_(*E*) = 0.32 *T, t*_*i*_(*M*) = 0.5 *T* and *t*_*i*_(*L*) = 0.68 *T* where *T* = 1 is the total movement duration. We estimated the model parameters of all individual participants from the actual experimental results – hand trajectories over trials – of the original 2-VP study [6]. These parameters describe the learning behavior of each player within a dyad.

The results of the identification procedure are summarized in Figure 4. To address how well the movement data fit the model, for each player we compared the actual and predicted actions. For both groups of players, we obtained an average *R*^2^ = 0.86 – see the Supplementary Note for details.

**Fig 4.**
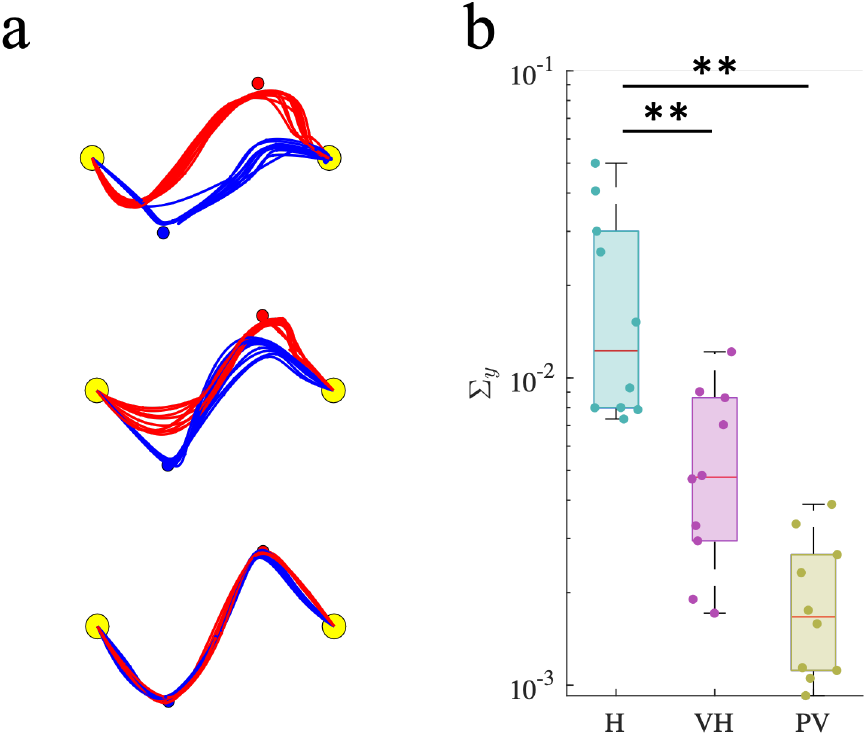
Fitting performances. a. Estimated trajectories in the identification procedure (last ten trials), for three representative dyads (one per group). b. Group differences in the estimated model parameters. The box plots display median, 25th and 75^th^ percentiles of the estimated parameters, for H (light blue), VH (pink) and PV (yellow) groups. Asterisks indicate statistically significant differences (*: p < 0.05; **: p < 0.01)

The predicted hand trajectories are consistent with the experimental findings – see Figure 4b – and exhibit distinct features in the different experimental conditions. We found significant group effects in the sensory variance parameter 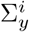 − F(2,27)=10.89, p< 0.001. Post-hoc analysis revealed that 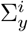 is significantly lower in the PV group respect to the H group (p< 0.001) and in VH with respect to the H group (p= 0.004). All the remaining parameters exhibited no significant group effects. Figure 4c summarizes these effects. The group differences in the estimated 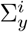 reflect the different experimental conditions – this suggests that the identification procedure has captured the latter from the movement data alone.

To further clarify the role of sensory noise in the development of joint coordination, we used the model to simulate the five dyads per each experimental group; we then compared the temporal evolution of the simulated movements in all three experimental conditions with the corresponding experimental results – see Figure 5. The simulated trajectories qualitatively resemble the experimental results – see Figure 5a. We specifically looked at the temporal evolution of inter-player coordination in terms of the minimum distances of the *i*-th player’s trajectory from the *j*-th via-point (MD_*ij*_, with *i* ≠ *j*) – see Figure 5b. Consistent with experimental findings [6], with lower sensory noise (PV group) the players converged toward a shared solution that is more similar – smaller MD_*ij*_ – to the theoretical Nash equilibrium - figure 3b. The simulated MD_*ij*_ time series for the three groups turned out to be very similar to the experimental observations. In particular, the rate of decrease was faster when the information about the partner was more reliable.

**Fig 5.**
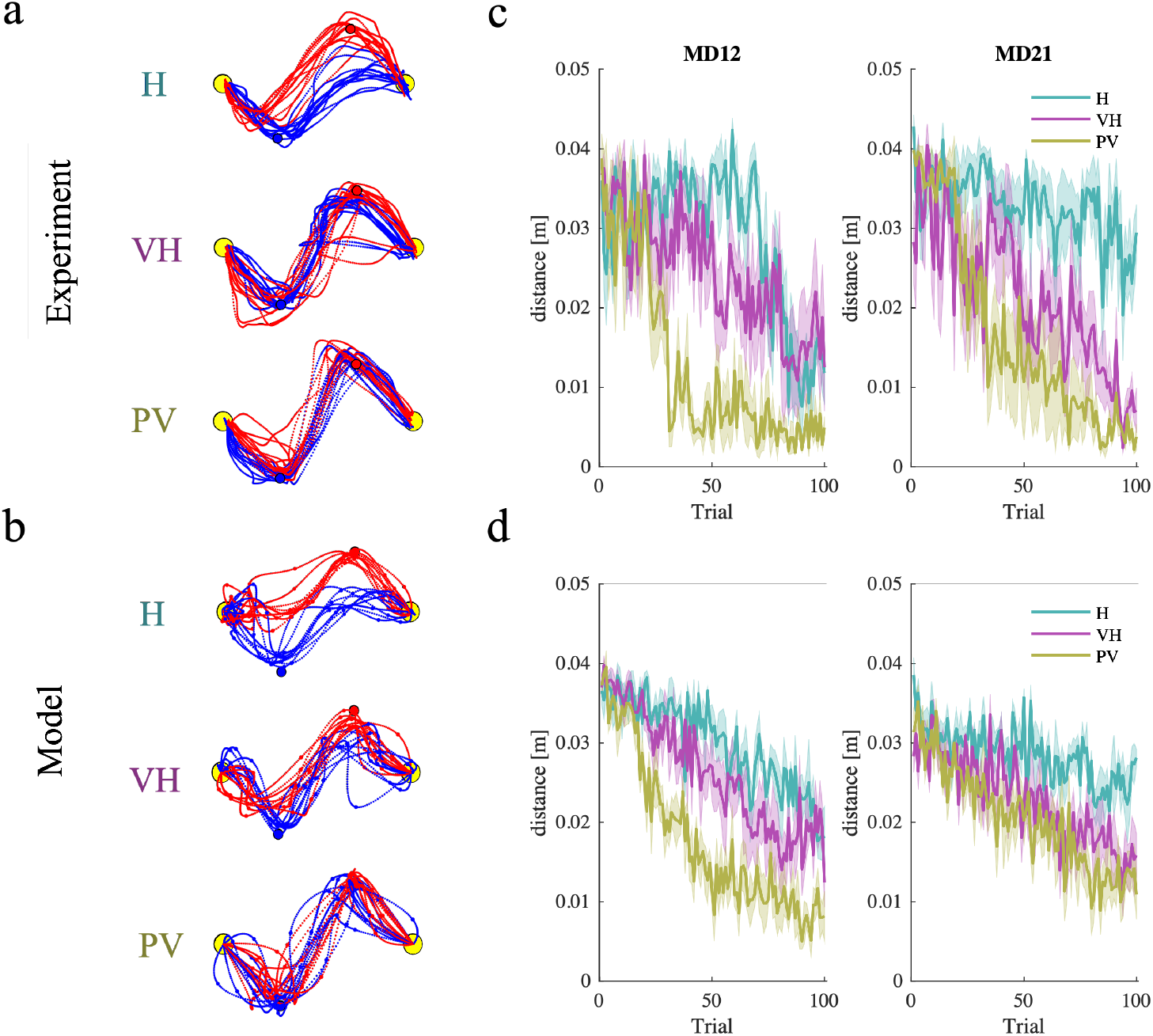
Experimental results (top) and simulations (bottom) of the 2VP task. a. Experimentally observed and b). simulated trajectories for selected dyads in the three experimental conditions (H, VH, PV) in the original 2-VP study [6]. c. Experimentally observed and d. simulated minimum distances of player 1 from VP_2_ (MD_12_) and player 2 from VP_1_ (MD_21_) – mean *±* SE over actual and simulated players – in all three conditions.

One of the main findings of the original 2-VP study [6] was that, as the information about the partner became less reliable, the players converged to a joint strategy that was more robust to partner uncertainty, in which the players actively controlled the crossing of their own via-point and passively followed the partner toward the other via-point, thus alternating leader and follower roles within the same movement. This effect was quantified in terms of a metric – leadership index, LI_*ij*_ – calculated as the average power of the interaction force of the *i*-th player just before crossing the *j*-th via-point. A negative or positive LI_*ij*_ indicates that the *i*-th player behaves, respectively, as a ‘leader’ or as a ‘follower’ – see the Methods section. The experimental findings point to a dependence of the observed leadership indices on the reliability of sensory information. To understand the nature of this relation, in the experimental data we compared the leadership indices calculated for each subject at the end of the training phase and the estimated sensory noise covariance 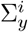 for that same player. In addition, we used the model to simulate dyads playing the game in which we systematically varied 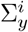, while keeping the other parameters constant. The relation between 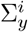 and the leadership indices is summarized in Figure 6a. In both experiments and simulations, when crossing their own via-pont both players behave as leaders (negative LI_11_ and LI_22_). Consistent with the simulations, the experimental data also suggest that in player 1 the amount of leadership (LI_12_) is significantly correlated to noise covariance (r= -0.632, p=0.011) whereas no such correlation is observed in player 2 (LI_21_). When considering the crossing of the partners’ via-points, both players exhibit a clear ‘follower’ behavior (positive LI_12_ and LI_21_), but only player 1 exhibits a significant correlation with the estimated noise covariance (r= 0.5799, p=0.023). Again, the simulations – Figure 6a, red and blue lines – are consistent with this observation.

**Fig 6.**
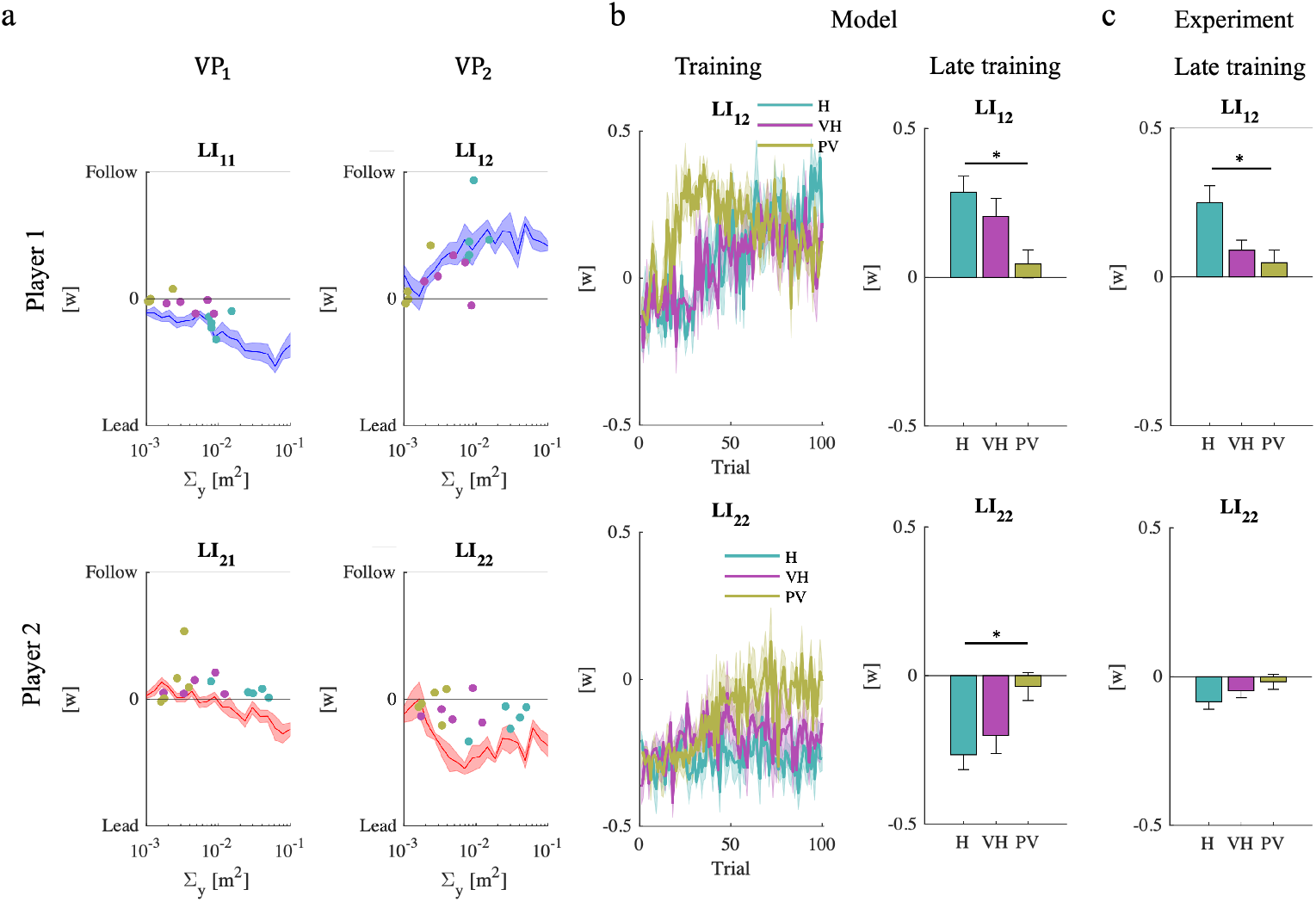
Sensory noise modulates the leadership index. a. Leadership indices as functions of the estimated sensory noise covariance 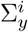 in experimental data (one data point per subject). 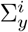 was systematically varied, and each 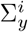 was simulated five times. The lines (mean *±* SE) denote the simulation results, for player 1 (blue) and player 2 (red). b. Simulated leadership indices over time (mean *±* SE); c. Simulated (left) and experimental (right) leadership indices for VP_2_ in the three experimental groups, at the end of the training phase. Experimental data from [6]

In summary, the experiments point at a stronger leader-follower role alternation in player 1 than in player 2. As a consequence, at VP_2_ the two players exhibit a much greater role differentiation: player 2 as leader – LI_22_ < 0 – and player 1 as follower – LI_12_ *>* 0). The model does not fully account for the different behaviors in VP_1_ and VP_2_ and other factors, possibly related to within-trial interaction, may play a role. Early in the movement, player 2 may focus on achieving their subgoal, thus being less prone to follow player 1. Conversely, later in the movement, player 1 – having reached their own subgoal – may be more willing to be led by player 2.

We then focused on the leadership indices for VP_2_. Figure 6b depicts the simulated temporal evolution of the leadership indices for player 1 and player 2 just before the crossing of the second via-point – respectively, LI_12_ and LI_22_. In the PV group, both indices quickly go to zero, indicating that roles tend to disappear as the interaction force disappears and the dyads get closer to the Nash equilibrium. In contrast, in the H and VH groups these indices change much more slowly and roles are preserved until the very end of the training phase. This model prediction is confirmed by statistical analysis of these quantities at the end of the training phase – significant group effect in LI_22_ (F(2,14) = 5.03, p = 0.0259) and in LI_12_ (F(2,14) = 5.13, p = 0.0245)– and is consistent with the experimental observations – see Figure 6c.

In conclusion, the model not only captures the consequences of varying the reliability of sensory information. Also, it correctly predicts the temporal evolution of the kinematics (minimum distances) and the kinetics (leadership indices) of the coordinated movements. Therefore, a simple best-response strategy seems to capture the ongoing development of joint coordination.

#### Experiment 2: symmetric via-points

When the via-points are arranged symmetrically - both VPs placed on the axis of the start-target line – the two Nash equilibria are cost-equivalent – see Figure 3 b. To develop stable coordination, the players face the additional challenge of converging to the same equilibrium. To investigate how this can be achieved we ran a novel study in which the experimental protocol was the same as in [6], but the via-points were arranged symmetrically. As in the previous study, in two experimental groups we manipulated the information available about the partner – either haptic alone (H) or visuo-haptic (VH).

Experiment 2 has an additional component – decision on strategy. To identify the strategy that the players followed at each trial, we used the minimum distance to the partner’s via-point (MD_*i*−*i*_) to determine whether that player got closer to their partner’s VP earlier or later in the movement – thus crossing or getting closer to the via-points in the VP_*i*_ → VP_−*i*_ order (E strategy for short); the VP_−*i*_ → VP_*i*_ order (L strategy); or they only focused on their own VP (M strategy); see the Methods section for details. The overall coordination performance is summarized in Figure 7.

**Fig 7.**
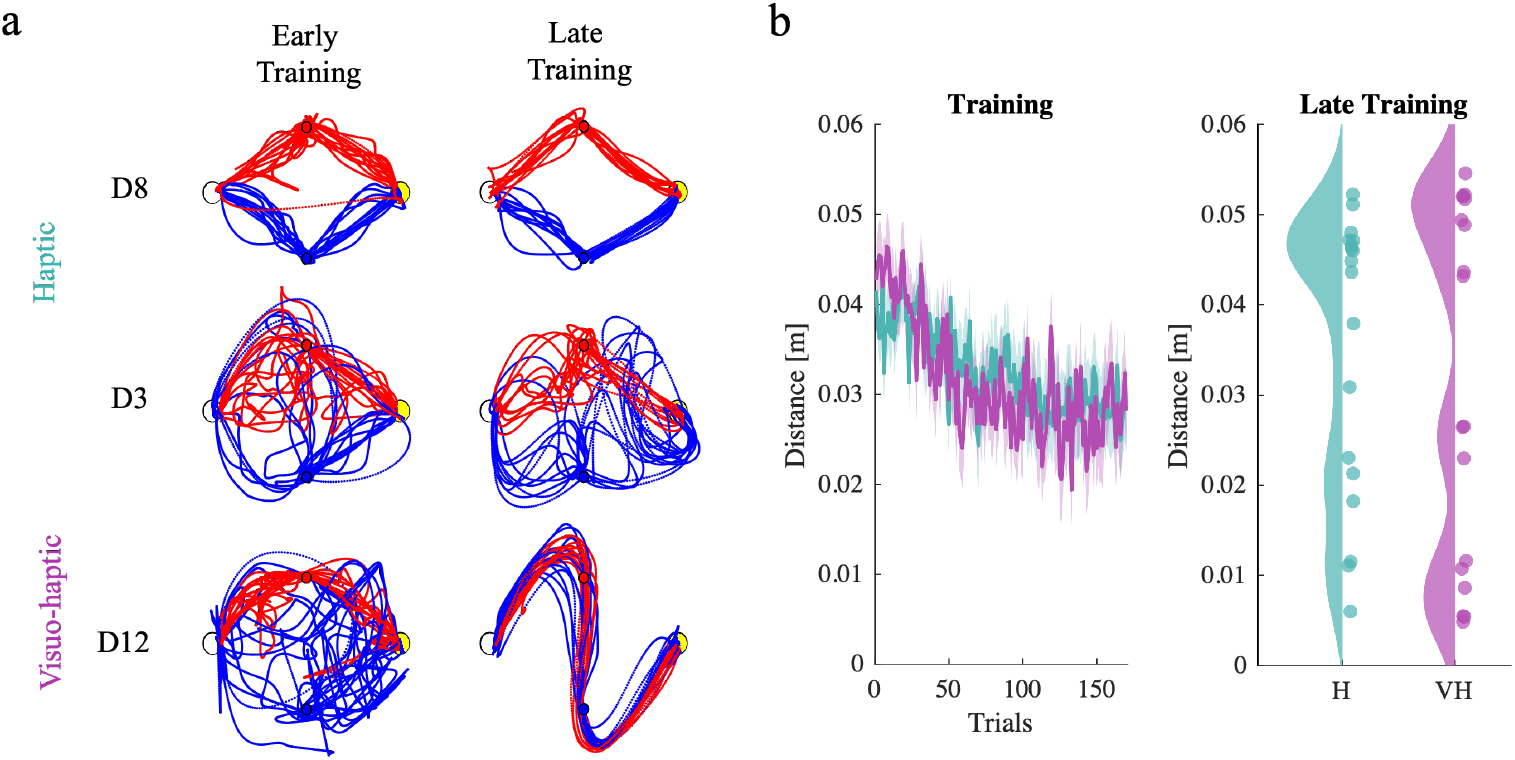
Coordination behavior in the symmetric 2-VP task. Dyads exhibit a variety of coordination behaviors. a. Typical dyad trajectories (from top to bottom: D8, D3, D12) at early and late training of the experiment. Player 1 and Player 2 are depicted in blue and red. These dyads exemplify typical behaviors: players ignoring each other (D8); cycling between opposite strategies (D3); and converging to a coordination (D12). b. Minimum distance of the two players from partner’s via-points. We report population average and standard error over time for the two groups. On the left we report distribution of the average performances at the end on training.

Joint coordination only occurs if the two players follow opposite strategies, i.e. either E-L or L-E. Figure 7a exemplifies these behavior types. In some dyads, one or both players gradually reduce their minimum distance from the opponent via-point – see Figure 7b.

In other dyads, the minimum distance does not decrease over training. Statistical analysis confirms an overall significant Time effect for both players (MD_12_: F(1,16) = 6.272, p = 0.0235; MD_21_: F(1,16) = 9.621, p = 0.0069), but there is neither a significant Group effect, nor a significant Time *×* Group interaction. However, a closer look at the MD_12_ and MD_21_ at the end of the training suggest that they have a bimodal distribution, indicating that in both experimental conditions only a fraction of the players attempt to incorporate partner’s action in their motor command. From this observation, we classified each player as a ‘Learner’ if their average MD_*ij*_ at the end of the training phase was below 0.02m; Non-Learner otherwise. Using this criterion, 7 out of 18 players were classified as Learners in the H group and 9 out of 18 in the VH group. The group difference is not statistically significant (*χ*^2^ test).

Overall, these findings suggest that forms of coordination that require the selection of one among multiple cost-equivalent solutions are much more complex processes than the asymmetric 2-VP settings [6], and coordination performance is not determined solely by the reliability of sensory information.

Nevertheless, a fraction of the dyads succeeded in achieving stable coordination in both space and time. To characterize individual behaviors, we estimated the model parameters from the sequence of the movements of each participant over trials. In the model, we set *t*_*i*_(*E*) = 0.38 *T, t*_*i*_(*M*) = 0.5 *T* and *t*_*i*_(*L*) = 0.62 *T* . We identified the model parameters for each player in a dyad from the time course of dyad movements. The fitting results are summarized in Figure8.

**Fig 8.**
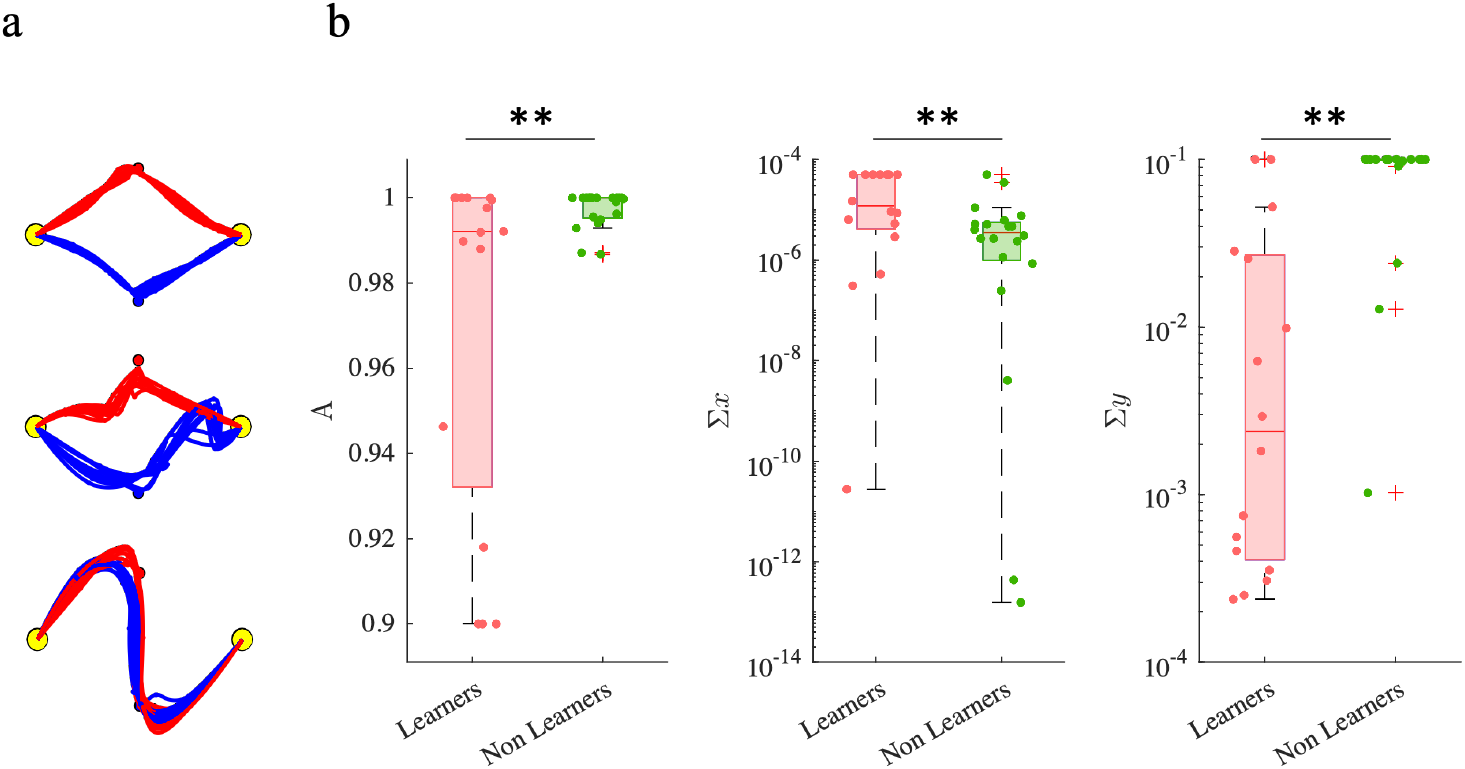
Determinants of learning. a. Estimated trajectories. b, c. Estimated model parameters grouped in terms of Learner and Non-Learner and in H and VH groups. Boxplots display median, 25th and 75th percentiles of the estimated parameters values, for Learners (orange) and Non-Learner (green). *: p < 0.05; **: p < 0.01.

Consistent with the experimental findings, no parameter exhibited a group (H, VH) effect (one-way ANOVA). We then examined differences in the Learner vs Non-Learner groups. In this case, three model parameters exhibited a Group effect. The sensory variance (∑_*y*_, is smaller in the Learner group – F(1,34) = 35.24, p < 0.0001. The internal process variance (∑_*x*_) is greater in the Learner group – F(1,34) = 8.43, p = 0.0065. The partner model retention rate (*A*) is smaller in the Learner group – F(1,34) = 8.5, p = 0.0062; see Figure 8b.

As in Experiment 1, we simulated the individual dyads. The simulations closely resemble the experimental findings. The model qualitatively predicts the different behaviors of Learners and Non Learners. The minimum distances from the partner via-points at the end of training did not significantly differ in the experiment and in the simulations in the Learners groups. We found a significant difference in the Non-Learners group (p = 0.0056) – see Figure 9a. In the experiments, Non-Learner dyads ignored their partner during the whole training phase. The model did not fully capture this behavior. This is possibly due to the fact that in these experiments the actions tend to be quite similar from trial to trial. The persistent excitation condition may not be satified, thus leading to inaccurate parameter estimates and ultimately to poor match between experiments and simulations.

**Fig 9.**
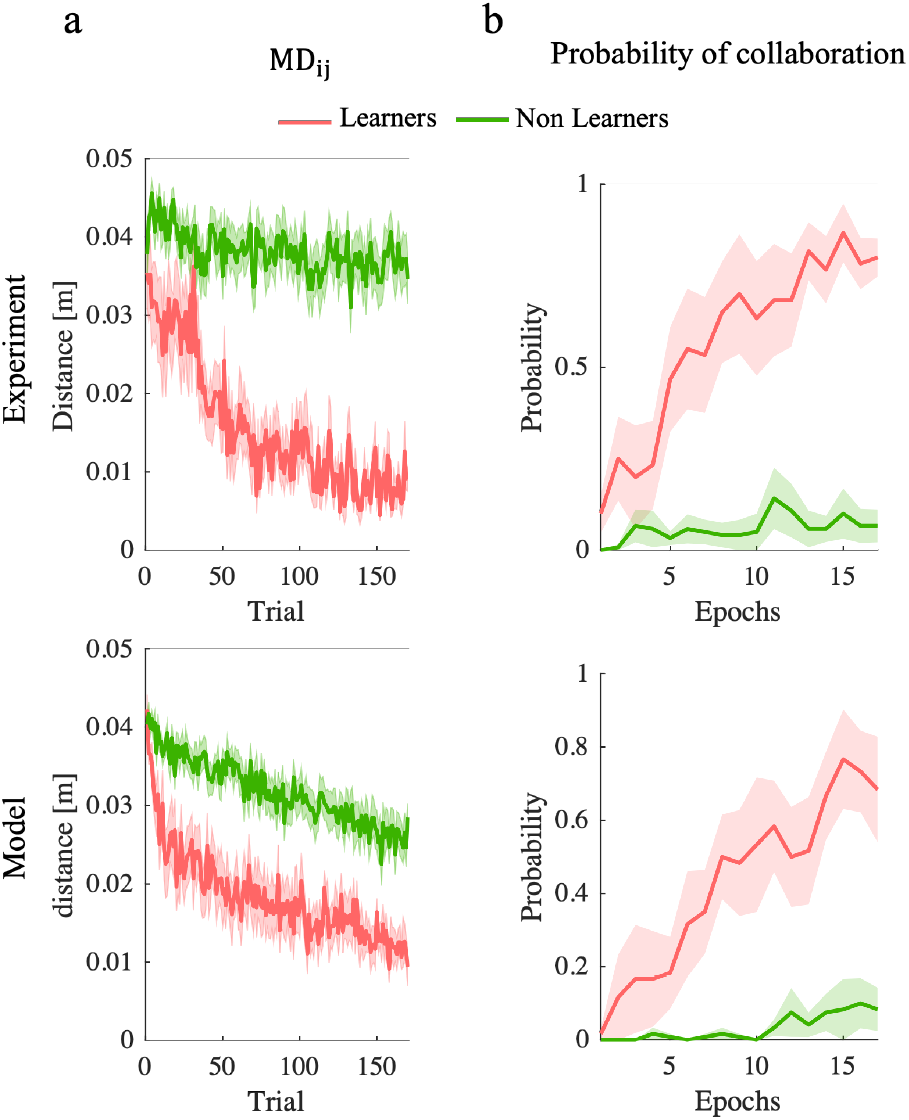
Simulations (bottom) closely resemble Experiments (top), in the temporal evolution of both the minimum distance to partner’s via-point over trials (a) and in the probability of establishing a collaboration (b), in both Learners (pink) and Non-Learners (green)

In simulated trials, the temporal evolution of the probability to establish a collaboration – see Figure 9b – is consistent with the experimental findings. In Learners, the probability tends to increase with time, whereas in Non-Learners the probability remains low.

## Discussion

We developed a general computational framework to describe how pairs of agents (dyads) gradually negotiate coordinated movements over multiple repetitions of the same interactive task. Each player is assumed to know their own goal; to have no information about their partner’s, and limited information about their action. At each trial, the *i*-th player selects an action (*u*_*i*_) on the basis of a partner model (*x*_*i*_). After both players have performed their actions, through their sensory system provides each player information (*y*_*i*_) about their partner’s action. This information is used to update the partner model to be used on the next trial. This modeling framework specifically focuses on how uncertainty about own task and uncertainty about the partner determine the game outcome.

The model can be used as a generative tool, to simulate joint action scenarios, or as an analytical tool to characterize individual players’ behaviors and strategies in actual joint action experiments. The simulation results may provide insights on the way coordination strategies develop. We demonstrated the model in the context of two different joint action scenarios: a motor version of the Stag Hunt game [27] and a joint 2-Via-Point task [6] with asymmetric and symmetric via-points. In the first scenario, we used simulations to explore the model’s parameter space in order to understand the determinants of emergent joint coordination. In the second scenario, we estimated the model parameters from experimental observations, and demonstrated that the model correctly captures the experimental manipulations and their effect on how joint coordination is learned.

### Trust as the basis of cooperation

In the Stag Hunt game, the players have the option to choose a safe but less profitable action – hunting a rabbit – which requires no coordination with their partners, or a potentially more profitable action – hunting a stag – which requires that both players aim at it to succeed. As they select their actions independently, under what conditions we can expect that they end up cooperating if they play the game repeatedly? If after each round both players are provided with imperfect information about their partner’s action, how is this information used to develop a propensity to behave cooperatively?

The Stag Hunt game has been used extensively as a model for the emergence of trust in social settings. Does it require an innate predisposition, or trust can be learned through repeated interaction? [28, 29]. Our model suggests an intermediate possibility. Under the assumption that own cost structure is completely known and thus there is no strategy learning, at each trial the prediction of partner actions optimally combines the sensory information available about the previous action, and the previous history of interaction. Hence the partner model adapts continuously over game iterations. The simulations indeed suggest that necessary conditions to develop ‘trust’ are (i) a prior belief that the partner does not behave erratically – in the model, this trait corresponds to a low 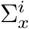; and (ii) reliable information about current partner action – low 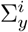; see Figure 2. In these situations, the Bayesian partner model has more ‘inertia’, in the sense that it places more weight on previous history of interaction and less on the last action. In its current implementation, partner representation is minimal, in the sense that only focuses on predicting the partner’s next action. More sophisticated partner representations, involving their goals and intentions, and possibly the way they represent their own (theory of mind) are likely even more effective in building up trust and thus cooperation [27].

### How is joint coordination learned?

In the asymmetric 2-VP task, model simulations replicated the gradual development of coordinated trajectories that was observed empirically [6] – see Figure5 and 6. In particular, the model correctly predicts that over trials, the hand paths undergo a gradual shaping process, in which they gradually reduce their distance from the partner’s via-point. Consistently, roles gradually disappear. Uncertainty about partner action modulates rates of convergence and final performances. This is remarkable as in the model the agents have no explicit information about their partner’s task requirements. The model also reproduces the three experimental conditions tested in the original study. Consistent with experimental observations, with lower sensory noise the players converged toward a shared solution that is closer to the theoretical Nash equilibrium. These results further support the notion that lower sensory noise facilitates the development of joint coordination.

### Achieving a joint coordination is not just a matter of more reliable sensory information

To further investigate how interacting agents negotiate a coordination over multiple game repetitions, we focused on a scenario in which the agents must select one among multiple strategies, and develop appropriate spatiotemporal coordination within that selected strategy. To do so, we manipulated the geometry of the 2-VP task so that the via-points were located symmetrically with respect to the horizontal axis, thus leading to two cost-equivalent Nash equlibria. This symmetric version of the 2-VP task is much more challenging than its asymmetric counterpart [6], because to achieve a collaboration, the players not only need to cross or approach their partner’s via-point; they must also decide whether to do so before or after crossing their own. Hence achieving a collaboration is more difficult, and may actually never occur.

In these experiments, we observed a gradual improvement of spatiotemporal performance which indicates that, at population level, the dyads increasingly coordinate their movements over trials. However, only a fraction of the dyads actually converged to a stable coordination. Some dyads got stuck in cyclic behaviors, in which both players switch between the two coordination solutions. In other dyads, each player only focused on their own VP by ignoring the mechanical interaction with their partner, even though this implied more mechanical effort on their side. This latter finding suggests that collaboration has a cost, which may overcome the extra mechanical effort required by lack of collaboration in this task. However, we found no evidence that convergence to stable coordination is determined by reliability of information about the partner. This finding is in partial contrast with previous observations [6], that more reliable sensory information has quantitative and qualitative effects on coordination, in the sense that the latter is faster and closer to the predictions of game theory (Nash equilibrium). Nevertheless, some dyads did converge to a shared strategy. The question arises of whether some specific sensorimotor abilities and/or peculiar features of partner representation facilitate this outcome.

The estimated model parameters did not capture the experimental manipulation of the sensory information (H and VH groups); this may be due to the fact the observed macroscopic behaviors – convergence or non-convergence to a stable coordination – are little different in the two experimental groups. It should be noted, however, that the relevant model parameter – sensory covariance 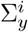 – reflects not only the experimental manipulation but also the individual sensory capability, which may vary in different individuals. However, when looking at Learners and Non-learners, we found that the former are characterized by a combination of lower sensory covariance 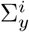, higher partner model covariance, 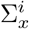 and lower partner model retention, *A*^*i*^. In other words, Learners are more prone to integrate current sensory feedback to predict their partner’s action, whereas Non Learners tend to keep their initial belief and fail to develop a stable coordination. Taken together, these features indicate that Non learners are less likely to modify their partner predictions and therefore to switch strategy from one trial to another. These findings suggest that not only sensory information has quantitative and qualitative effects on coordination [6] but also some other specific sensorimotor abilities and the peculiar aspects of partner representation facilitate this outcome.

### Relation to other models of joint action

Previous models of joint action were purely theoretic [27, 30] or were only capable of reproducing qualitative aspects of the observed behaviors [5, 6]. The proposed model extends approaches used to model sensorimotor adaptation and relies on optimality principles to model both the estimation of partner actions and the selection of own action.

The model can be related to the active inference framework [31]. Active inference aims at unifying perception and action selection in terms of the minimization of one single quantity – surprise, or the mismatch between observation and predictions. Previous studies have addressed specific aspect of joint action on an active inference perspective – namely, theory of mind [27], or discriminating between self and joint action [32]. Recently, active inference has been proposed as a possible general framework to study joint action [33] and to explain the collective behaviors of multiple interacting agents [34]. Very much like active inference, we model perception of the partner as Bayesian estimation – by optimally combining observations and model predictions (in our case, of partner’s action) and by updating the model in terms of the prediction error. In our model, each player selects the action that minimizes their expected cost, which combines a task representation – for instance, an intended target – that is supposed to be known – and the partner model. In the active inference framework the action cost may be seen as playing the role of a prior [35] for action selection. In our model the two processes – (partner) perception and action selection are kept separate, which allows to use the tools of game theory to address the development of joint coordination.

### Model limitations and possible extensions

The model can be applied to different interactive settings. We addressed a setting where the matrices defining the cost function are positive definite, then the conditional action probability is a Gaussian distribution. Otherwise - as in competitive games - the action probability may not be Gaussian although the model formulation is still valid. The only change is in the system identification procedure.

In the presented formulation we made a series of simplifying assumptions. However, the model must be considered as a basic framework, which can be easily extended to more complex scenarios. First, the model assumes that each players has a perfect knowledge of their own goal – the structure of the cost function – and no knowledge at all about their partner’s. Decay of the action selection temperature *λ*_*i*_(*t*) can be interpreted as a gradual refinement of the cost function, but it is possible to add to the model a reinforcement learning mechanism in which a reward – function of the cost – provided to each player at the end of each trial – can be used to update the cost function parameters. Second, the model uses fictitious play to model the development of coordination. Fictitious play assumes that the players select their action on the basis of information about their partner’s last action or, more in general, a probabilistic model of their partner’s actions, under the assumption of a stationary strategy. Our simulations suggest that this model produces plausible behaviors and – at least in two studies [4, 6] – reproduces well the reported joint action time series. However, this does not rule out the possibility that in other joint action scenarios, the players also develop a model of their partner’s goals – encoded in the structure of their cost function. This information could be used, for instance, as a prior in the partner model [36]. The players may use even more complicated insight - e.g., the partner’s mental state or ‘theory of mind’ to make their decisions. The partner’s mental state can be simply made part of the partner model, can be inferred from observation of the action outcomes, and accounted for in the expected cost during action selection.

## Conclusions

In summary, our proposed model successfully captures the experimental findings across various strategic scenarios. The model provided valuable predictive capabilities and highlighted the importance of reliable information and exploration in achieving Nash equilibria. The parameter identification process further elucidated the impact of experimental conditions on the estimated model parameters. Overall, our model serves as a useful tool for understanding and predicting strategic decision-making processes in diverse settings.

Overall, these findings confirm that model-based analysis of joint action experiments may provide valuable information on the underlying mechanisms. In particular, we addressed joint action using a pair of computational models representing the two interacting agents. Coupled computational models that evolve over time to describe the structure and behavior of a physical system or process are referred to as digital twins. Digital twins and computational models of joint action are generally complicated frameworks which require custom implementations. The presented model relies on dynamical system theory and probabilistic graphical models. This is consistent with previous works [37] proposed a general and robust foundation for the development of digital twins. As such, the model described in this chapter is a promising tool to advance the understanding of the processes underlying joint action and a wide variety of applications can be envisioned. In healthcare, computational models or digital twins of interacting agents promise to advance medical assessment, diagnosis, and personalized treatment. In a similar way, in educational settings, they can be used to devise personalized training paths.

## Methods

### Spatial Stag Hunt

The parameters of the cost function can be calculated in order to achieve the cost matrix reported in Figure 2. In particular, in Eq. 12 we set *J*_1_(*u*_*R*_, *u*_*R*_, 1) = *J*_1_(*u*_*R*_, *u*_*S*_, 1) = 5, *J*_1_(*u*_*S*_, *u*_*R*_, 0) = 10, and *J*_1_(*u*_*S*_, *u*_*S*_, 0) = 1 – same for *J*_2_. This specifies three equations, which uniquely define the three parameters: *w* = 9/2, *z*_*R*_ = 5, and *z*_*S*_ = 1. The model parameters were finally set as follows:

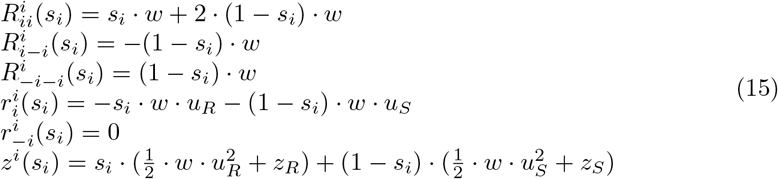

We simulated this game scenario under two different levels of sensory noise variance 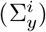 (low: 0.1; high: 10) and player’s belief on partner predictability 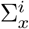 (high predictability: 0.1; low predictability: 10). For all other model parameters we set the following baseline values: *μ*^*i*^=0, 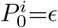, *A*^*i*^=0.99, 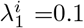 and *a*^*i*^=0.999. For each combination of 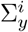 and 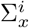 we simulated eight dyads. For each player in a dyad we applied random variations (within a *±* 10 % range) to the baseline parameter values. We repeated the game a total of 2000 times (50 blocks of 40 trials each). We used a 2-way ANOVA to assess the effect of the two factors (sensory noise: low-high and internal noise: low-high).

### Two Via-Point Task

#### Experiments

The experimental protocol is identical to [6]. Each participant sat in front of a computer screen and grasped the handle of a haptic interface (Novint Falcon). They could not see or hear each other and were not allowed to talk. The participants were instructed to perform planar point-to-point movements in the vertical plane, between the same start and end point (displayed, respectively as a white and a yellow circle of ⊘ 1 cm), but through different via-points (a blue circle for Player 1, a red circle for Player 2, both with ⊘ 0.5 cm. In a reference frame centered on the robot workspace, with the X axis aligned with the left-right direction and the Y axis aligned with the vertical direction, for both players the start point was placed in the (-5, 0, 0) cm position and the target point was placed in the (5, 0, 0) cm position. Hence the start and the target point had a horizontal distance of 10 cm. In Experiment 1 the via-points were placed, respectively, at locations VP_1_ = (-2,-3,0) cm and VP_2_ = (2,3,0) cm, asymmetric with respect to the Y axis. In Experiment 2 the via-points were placed, respectively, at locations VP_1_ = (0,-3,0) cm and VP_2_ = (0,3,0) cm, symmetric with respect to the Y axis. The two participants were mechanically connected through the haptic interfaces, which generated a force proportional to the difference of the two hand positions, i.e. *F*_*i*_ = −*k*(*x*_*i*_ − *x*_−*i*_). The experimental protocol was organized into epochs of 12 movements each and consisted of three phases: (i) baseline (1 epoch), (ii) training (17 epochs), and (iii) wash-out (2 epochs) for a total of 20 *×* 12 = 240 movements. During the baseline phase, the interaction forces were turned off and each subject performed on their own (‘solo’ performance). During the interaction phase, the participants were told that they might experience a force while performing the task and were instructed to keep this force to a minimum. We introduced this additional requirement because of a technical limitation of our robots, which are unable to generate large stiffnesses. With a low stiffness the haptic perception of the interaction force is less reliable, and introducing an explicit requirement on low interaction forces made sure that all players were provided with exactly the same task requirements irrespective of the information they had available about their partner.

In Experiment 1 we reanalyze the data from [6] – a total of 30 participants forming 15 dyads, matched by their body mass index (BMI). Each dyad was randomly assigned to three groups (5 dyads per group), which differed in the available sensory information. In the haptic (H) group the interaction could only be sensed haptically. In the visuo-haptic (VH) group the interaction force (magnitude, direction) was also displayed as an arrow attached to the cursor (scale factor: 10 N/cm). In the partner-visible group (PV), in addition to the interaction force the partner position was continuously displayed on the screen. In Experiment 2, we recruited 36 participants, all right handed (Edinburgh Handedness Inventory [38]). All participants were naive to the task and had no known history of neurological or upper limb motor impairment. We formed 18 BMI-matched dyads. The two participants within the same dyad were randomly labeled as, respectively, Player 1 and Player 2. Each dyad was randomly assigned to two groups (H: 6 M +12 F, age 25 *±* 3; VH: 8 M + 10 F, age 24 *±* 5), with the same meaning as Experiment 1. The study conforms to the ethical standards laid down in the 1964 Declaration of Helsinki that protects research subjects and was approved by the Institutional Review Board (Comitato Etico Regione Liguria). All participants provided written informed consent to participate in the study.

#### Data analysis

During experiments, hand trajectories and the interaction forces generated by the robots were sampled at 100 Hz and stored for subsequent analysis. Movement and force trajectories were smoothed using a 4th order Savitzky-Golay filter with a 370 ms time window. The same filter was used to estimate velocity and acceleration. A sign of coordination is that each player, while passing through their own via-point, also gets very close to their partner’s. This can be quantified in terms of the Minimum Via-Point Distance, MD_*ij*_ of player *i* from via-point *j*: MD_*ij*_ = min_*t*_ dist_*ij*_(*t*), with dist_*ij*_(*t*) = ||*x*_*i*_(*t*) − *x*_VP*j*_||. A pre-requisite for coordination is that both players cross their partner’s via-point, i.e. both reduce their MD_*i*−*i*_. We classified each trial as collaborative in space if both players had MD_*ij*_ < *d*, with *d* = 0.02 m. The threshold was set based on the observation of dyads who clearly exhibited a coordination. We then estimated the decision variable *s*_*i*_(*t*) in which at every trial *t* player *i* selects his/her action *s*_*i*_(*t*) among three options: moving to VP_*i*_ and then VP_−*i*_ (E strategy); moving to VP_−*i*_ and then VP_*i*_ (L strategy); or moving to own VP while ignoring the other (M strategy). In summary:

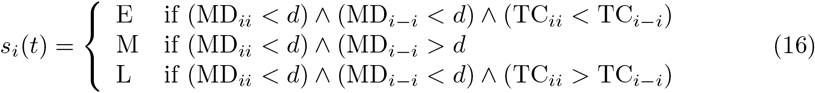

where TC_*ij*_ is the time instant – with respect to the dyad start time – at which player *i* achieves its minimum distance to via-point *j*: TC_*ij*_ = arg min_*t*_ dist_*ij*_(*t*); therefore, MD_*ij*_ = dist_*ij*_(TC_*ij*_). In order to understand the determinants of learning, we defined a player as a ‘learner’ if he/she consistently achieved either E or L during the last epoch of the training phase; ‘non-learner’ otherwise. Note that both players in a dyad being learners according to this definition does not necessarily imply that they achieve a collaboration – they may get stuck into a cyclic behavior in which they alternate E and L strategies and never converge. We then focused on dyad behaviors. We classified a dyad as collaborative if both players on that trial used either E-L or L-E:

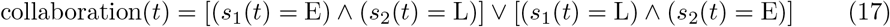

In a similar way, we classified a trial as ‘cyclic’ if the two players took either E-E or L-L, and as ‘non-collaborative’ if at least one of the players ignores the other. Using these definitions, for each dyad we calculated the probability of collaboration, cycling and non-collaboration over each 12-trial epoch. We also calculated a leadership index (LI_*ij*_) – the power of the interaction force of the *i*-th player, averaged over the 300 ms interval just before crossing the *j*-th via-point. A negative or positive LI_*ij*_ denotes, respectively, that the *i*-th player moves against the interaction force – thus behaving as a ‘leader’ – or just passively complies with it – a ‘follower’.

For all indicators, we ran a repeated-measures ANOVA with group (H, VH, PV or V, VH) and Time (early – training epoch 1 – and late – epoch 17) as factors. We used post-hoc analysis (Tukey’s HSD) to further identify significant differences between group pairs in the case of significant group effects.

**Model** We approximated the trajectory at time *t* as a polyharmonic spline:

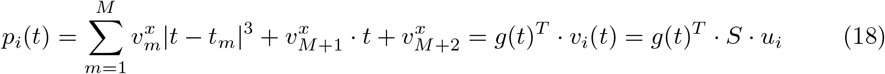

where *g*(*t*) = [|*t* − *t*_1_|^3^ … |*t* − *t*_*M*_ |^3^ *t* 1]^*T*^ and *S* is a constant matrix. Considering that some components of 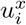 (and 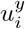) are fixed (initial and final position), we finally have

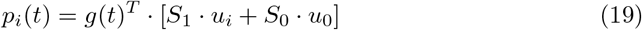

We defined the ‘action’ of the *i*-th player at a given trial as the vector *u*_*i*_ of nodes’ coordinates. This approximation captures the overall shape of a generic trajectory in terms of the coordinates of a small number of nodes. Subsequent derivatives of *p*_*i*_(*t*) (velocity, acceleration) can be also expressed as linear combinations of the *u*_*i*_’s. In all simulations, we used *M* = 10. In this way, Eq.13 becomes a quadratic form in *u*_*i*_ and *u*_−*i*_. Parameters *w*_1_, *w*_2_, and *r* were determined by the Bryson rule: 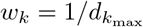 where 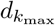 is the maximum acceptable magnitude of the corresponding term of the cost function. In our task, we set 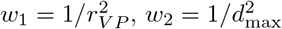 and 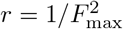, where the VP radius is *r*_*V P*_ = 2.5 mm, the maximum between-subject distance is *d*_max_ = 2 mm and the maximum force is *F*_max_ = 1 N.

For both experiments, we estimated the model parameters for all individual participants from the actual experimental data – movement sequences over the 100-trial training phase. These parameters describe the learning behavior of each player within their dyad. For each model parameter, we used a one-way ANOVA with Group (Experiment 1: H, VH, PV; Experiment 2: H, VH) as factor. In the case of significant group effects, we used post-hoc analysis (Tukey’s HSD) to further identify significant differences between group pairs.

To further clarify the role of sensory noise in the development of joint coordination, we used the model as a generative tool, by simulating the temporal evolution of the movements in all experimental conditions. To simulate individual dyads, we used the exact same parameters estimated in the fitting procedure for that dyad.

## Acknowledgments

The work is supported by project Fit for Medical Robotics (Fit4MedRob), National Plan for Complementary Investments (PNC) - National Plan for Recovery and Resilience (NPRR), CUP : B53C22006950001.

